# Tinnitus development is associated with synaptopathy of inner hair cells in Mongolian gerbils

**DOI:** 10.1101/304576

**Authors:** Konstantin Tziridis, Jan Forster, Isabelle Buchheidt-Dörfler, Patrick Krauss, Achim Schilling, Olaf Wendler, Elisabeth Sterna, Holger Schulze

**Affiliations:** Experimental Otolaryngology, Department of Otorhinolaryngology, Head & Neck Surgery, Friedrich Alexander University Erlangen-Nürnberg (FAU), 91054 Erlangen, Germany; Department of Otorhinolaryngology, Head & Neck Surgery, Friedrich Alexander University Erlangen-Nürnberg (FAU), 91054 Erlangen, Germany

**Author notes:** Corresponding Author: Dr. Konstantin Tziridis, Experimental Otolaryngology, Department of Otorhinolaryngology, Head & Neck Surgery, Friedrich Alexander University Erlangen-Nürnberg (FAU), Waldstrasse 1, 91054 Erlangen, Germany.

**Keywords:** auditory brainstem response, gap prepulse inhibition of the auditory startle response, rodents, cochlea, hearing loss

## Abstract

Human hearing loss (HL) is often accompanied by comorbidities like tinnitus which is affecting up to 15% of the adult population. Rodent animal studies could show that tinnitus may not only be a result of apparent HL due to cochlear hair cell damage but can also be a consequence of synaptopathy at the inner hair cells (IHC) already induced by moderate sound traumata. Here we investigate synaptopathy previously shown in mice in our animal model, the Mongolian gerbil, and relate it to behavioral signs of tinnitus. Tinnitus was induced by a mild monaural acoustic trauma leading to monaural noise induced HL in the animals, quantified by auditory brainstem response (ABR) audiometry. Behavioral signs of tinnitus percepts were detected by measurement of prepulse inhibition of the acoustic startle response in a gap-noise paradigm. 14 days after trauma, the cochleae of both ears were isolated and IHC synapses were counted within several spectral regions of the cochlea. Behavioral signs of tinnitus were only found in animals with IHC synaptopathy, independent of type of HL. On the other hand, animals with apparent HL but without behavioral signs of tinnitus showed a reduction in amplitudes of ABR waves I&II but no significant changes in the number of synapses at the IHC. We conclude – in line with the literature – that HL is caused by damage to the IHC or by other reasons but that the development of tinnitus, at least in our animal model, is closely linked to synaptopathy at the IHC.

## Introduction

Around 10 to 15% of the general adult population suffer from subjective tinnitus of some kind (Heller, 2003). This phantom sound condition can lead to insomnia, psychological disorders or, for the most severe cases, even suicide (Coles, 1984; Lewis et al., 1994; Langguth et al., 2011). The exact mechanism and the origin of this maladaptive process within the auditory pathway are still under debate (Gerken, 1996; Eggermont, 2003; Eggermont & Roberts, 2004; Engineer et al., 2011; Knipper et al., 2011; Schaette & McAlpine, 2011; Ruttiger et al., 2013), but most researchers consider malfunction of the inner ear – caused by presbyacusis, ototoxic pharmaceutics, injuries or noise traumata – to be the primary cause for the development of central tinnitus. On the other hand, also patients and animals without a measurable hearing loss can experience such an auditory misperception, indicating also a role of neuropathic deafferentation in its development (Weisz et al., 2006; Savastano, 2008; Hickox & Liberman, 2014). In this study we want to focus on noise induced tinnitus with or without a measurable hearing loss in an animal model of a possible pure tone or narrow band tinnitus percept.

Currently, it is still unknown why some patients with or without hearing loss do develop chronic tinnitus while others with comparable psychophysical or audiometrical assessed hearing sensitivity do not, although first animal results and models have been published trying to solve this riddle (Knipper et al., 2011; Schaette & McAlpine, 2011; Ahlf et al., 2012; Ruttiger et al., 2013; Tziridis et al., 2015). In mice it has been shown that a relatively mild acoustic trauma (8–16 kHz octave-band noise at 98 dB SPL for 2 h) may lead to a massive loss of synapses (= synaptopathy) at the inner hair cells (IHC) forming the afferent output of the cochlea, and subsequently to a deafferentation (= neuropathy) of the receptor epithelium (Liberman et al., 2015). This synaptic loss recently has been termed hidden hearing loss (Schaette & McAlpine, 2011), and has been investigated in the context of tinnitus in mice (Hickox & Liberman, 2014) and rats (Ruttiger et al., 2013). In their study, although conducted with a relatively low spectral resolution comprising just the two frequencies 11.3 kHz and 32 kHz, Hickox and Libermann found tinnitus percepts – based on the change of the prepulse inhibition (PPI) of the acoustic startle response – only in normal hearing but neuropathic animals but not in those without neuropathy. Consistently, Rüttiger and colleagues were able to show a significant reduction of IHC synapses after a mild mid-frequency trauma (120 dB SPL at 10 kHz for 1 h) in high-frequency regions of the cochlea only in animals that subsequently developed a tinnitus percept (based on foraging behavior for sugar water during sound presentation) but did not report a hearing loss in these animals.

In this report we investigated mild pure tone trauma (115 dB SPL at 2 kHz for 75 min) induced synaptopathy at IHCs and auditory brainstem response (ABR) wave changes with high spectral resolution (1 to 16 kHz in octave steps) in Mongolian gerbils with or without behavioral signs of tinnitus, as assessed by PPI of the acoustic startle reflex (ASR) (Turner et al., 2006; Turner, 2007; Ahlf et al., 2012), and with or without permanent noise induced hearing loss as assessed by auditory brainstem response (ABR) audiometry. In our animal model we found a clear link between tinnitus development and synaptopathy at the IHCs but not between tinnitus development and hearing loss. We speculate that the hearing loss may be related to – among several other potential causes –an additional outer hair cell (OHC) damage (Salvi et al., 2013).

## Methods

### Ethical statement

Mongolian gerbils (*Meriones unguiculatus*) were housed in standard animal racks (Bio A.S. Vent Light, Ehret Labor-und Pharmatechnik, Emmendingen, Germany) in groups of 2 to 3 animals with free access to water and food at 20 to 24°C room temperature under 12/12 h dark/light cycle. The use and care of animals was approved by the state of Bavaria (Regierungspräsidium Mittelfranken, Ansbach, Germany, No. 54-2532.1-02/13). A total of 19 ten to twelve weeks old male gerbils from Charles River Laboratories Inc. (Sulzfeld, Germany) were used in this study.

### Behavioral testing and auditory brainstem responses (ABR)

All animals were handled before the beginning of the experiments and accustomed to the setup environment to minimize stress. All behavioral and electrophysiological methods used have previously been described in detail (Ahlf *et al*., 2012; Tziridis *et al*., 2012; Tziridis *et al*., 2015; Schilling *et al*., 2017) and therefore are only briefly summarized here: During the first week, animals were tested first, with a GAP-NOGAP paradigm of PPI of the ASR (= gap-prepulse inhibition of the acoustic startle reflex, GPIAS) (Turner *et al*., 2006; Schilling *et al*., 2017) to acquire baseline data for later tinnitus assessment. 60 stimuli of band-pass filtered background noise (60 dB SPL) with center frequencies of 1 kHz, 2 kHz, 4 kHz, 8 kHz, and 16 kHz and a half octave width were presented with (30 stimuli) or without (30 stimuli) a 20 ms gap 100 ms before a startle stimulus (white noise burst, 20 ms duration, 2 ms sin^2^-ramp, 105 dB SPL). A complete behavioral measurement of 300 stimuli had a rough duration of one hour. Second, frequency-specific ABR measurements of both ears were performed under ketamine-xylazine anesthesia (0.3 ml/h: ketamine 500 mg/kg, xylazine 25 mg/kg) to obtain individual baseline audiograms for stimulation frequencies between 1 and 16 kHz in octave steps for stimulation intensities ranging from 0 to 90 dB SPL in 5 dB steps. For each ear, stimulus, and intensity, 300 repetitions were presented. The complete measurement of one ear took around 30 min. For the measurements three silver electrodes were placed subcutaneously retroaural above the bulla of the tested ear (recording electrode), central between both ears (reference electrode) and at the basis of the tail (ground electrode). The signal was recorded and filtered (bandpass filter 400 to 2000 Hz) via a Neuroamp 401 amplifier (JHM, Mainaschaff, Germany). One week after these measurements, a mild acoustic pure tone trauma (2 kHz, 115 dB SPL, 75 min) was induced monaurally under deep ketamine-xylazine anesthesia via a Canton Plus X Series 2 speaker placed directly in front of the traumatized ear in 15/19 animals; the non-traumatized control ear was protected using earplug foam (3M™ earplugs 1110, 3M, Neuss, Germany) adding at least 20 dB attenuation in the given frequency range (Stuermer & Scheich, 2000). In 4/19 animals a sham trauma was performed with the same parameters as above but only 65 dB SPL sound intensity. The extent of the trauma or sham was evaluated by a second binaural ABR measurement 3 to 5 days after the treatment and used to group the 15 traumatized animals into those with a significant, subacute mean hearing loss of at least 10 dB at the traumatized ear (10/15 animals; *apparent hearing loss group*) and those without (5/15 animals; *hidden hearing loss group;* as confirmed by later analysis of synaptopathy of IHC, cf. Histology below). In all 19 animals a possible persistent tinnitus percept was identified by a second GPIAS of acoustic startle response measurement and calculation of the GPIAS change (Schilling *et al*., 2017) 13 days after the trauma.

### Histology

14 days post trauma, after obtaining all behavioral and electrophysiological data, the animals were sacrificed, and the cochleae of the traumatized and non-traumatized ears were extracted and fixed in 4% formaldehyde for 1 hour. After de-calcification in 0.1 M EDTA for 1-2 days, the cochlear turns were usually cut into 3 pieces and immunostained for synaptic ribbon protein carboxy-terminal binding protein 2 (CTBP2; Khimich *et al*., 2005). Cochlear whole mounts were immunostained overnight at 4°C with primary antibody against CTBP2 (mouse anti-CTBP2 at 1:200; BD Transduction Labs, Heidelberg, Germany). Secondary antibody (donkey anti mouse conjugated with Cy3, 1:400, Dianova, Hamburg, Germany) was applied for 1h at room temperature.

### Data evaluation and statistical analysis

The obtained data of the GPIAS and ABR measurements were evaluated objectively and automatically by custom made Matlab programs (MathWorks, Natick, MA, USA). Tinnitus development was tested for each animal and frequency individually by t-tests of the log-normalized PPI: The log-normalization of the response amplitudes of gap (A_gap_) and no-gap (A_no-gap_) by log(A_gap_ / A_no-gap_) is necessary as it has been shown that only after this calculation parametrical testing (e.g. by t-tests) is allowed (Schilling *et al*., 2017). A significant decrease (p<0.05; Bonferroni corrected) of the GPIAS-induced change of the response to the ASR after the trauma (GPIAS_post_) relative to conditions before the trauma (GPIAS_pre_) in that test was rated as an indication of a tinnitus percept at that specific frequency. With this behavioral approach we are able to group animals independent of any other measurement into those with behavioral signs of a tinnitus percept and those without (Schilling *et al*., 2017).

The threshold determination was performed by an automated objective approach using the root mean square (RMS) values of the ABR amplitudes fitted with a hard-sigmoid function utilizing the background activity as offset (Schilling *et al*., 2019). The mean threshold was set at the level of slope change of this hard-sigmoid fit independent for each frequency and for each time point. Hearing loss was calculated by subtracting the post from the pre trauma thresholds and tested by parametric tests.

Peak-to-peak amplitudes of the ABR waves I&II, III, IV and V – corresponding to the activity in the cochlear nerve/dorsal cochlear nucleus, inferior olivary complex/trapezoid body, lateral lemniscus and inferior colliculus (Henry, 1979), respectively – were extracted from measurements at 65 dB SPL pre and post trauma from both ears. Where detection of earlier waves was not possible, later waves were identified according to their known latencies (cf. Tziridis *et al*., 2015). Mean ABR wave amplitudes were then analyzed using 2-factorial ANOVAs. Additionally, from the 90 dB SPL recordings (valid wave detections in all cases) the RMS values were calculated within a time window from 1 to 7 ms after stimulus onset from the mean ABR signal of each tested frequency, realigned for its distance to the behaviorally determined tinnitus frequency (in oct) and analyzed by 2-factorial ANOVAs.

The number of synapses per IHC was counted from the microscopic images (cf. below) by eye by one investigator without knowledge of the status of the cochlea (traumatized / non-traumatized ear) or any other attribute of the animal: The frequency map of the cochlea from 500 Hz to 16 kHz in half octave steps (Figure 1A, left panel) was created with the software Keyence BZ-II-Analyzer by measuring the relative distances beginning from the apex of the cochlea along the spiral, according to the published cochlear frequency map for the Mongolian gerbil (Müller, 1996). Full-focus microscopic images in the range of the synapses of the IHCs of the resulting 11 regions from the subjacent inner spiral bundle to the nerve terminal in the supra-nuclear region including all visible synaptic ribbons were obtained using a fluorescent microscope BZ9000 (Keyence, Neu-Isenburg, Germany) with a 40x objective (0.6 numerical aperture) and visualized using a ImageJ plugin. Labeled synapses of the IHC region were counted for groups of 15 IHC (Figure 1A, right panel), according to the matching regions of the frequency map. The obtained number of synapses was evaluated separately for tinnitus and non-tinnitus groups, as well as for trauma and control side counts (Figure 1B). Subsequently, all statistical analyses of the synapse counts were performed by nonparametric Kruskal-Wallis ANOVAs in Statistica 8 (StatSoft, Hamburg, Germany) as one cannot expect synapse counts to be normally distributed.

**Figure 1.**
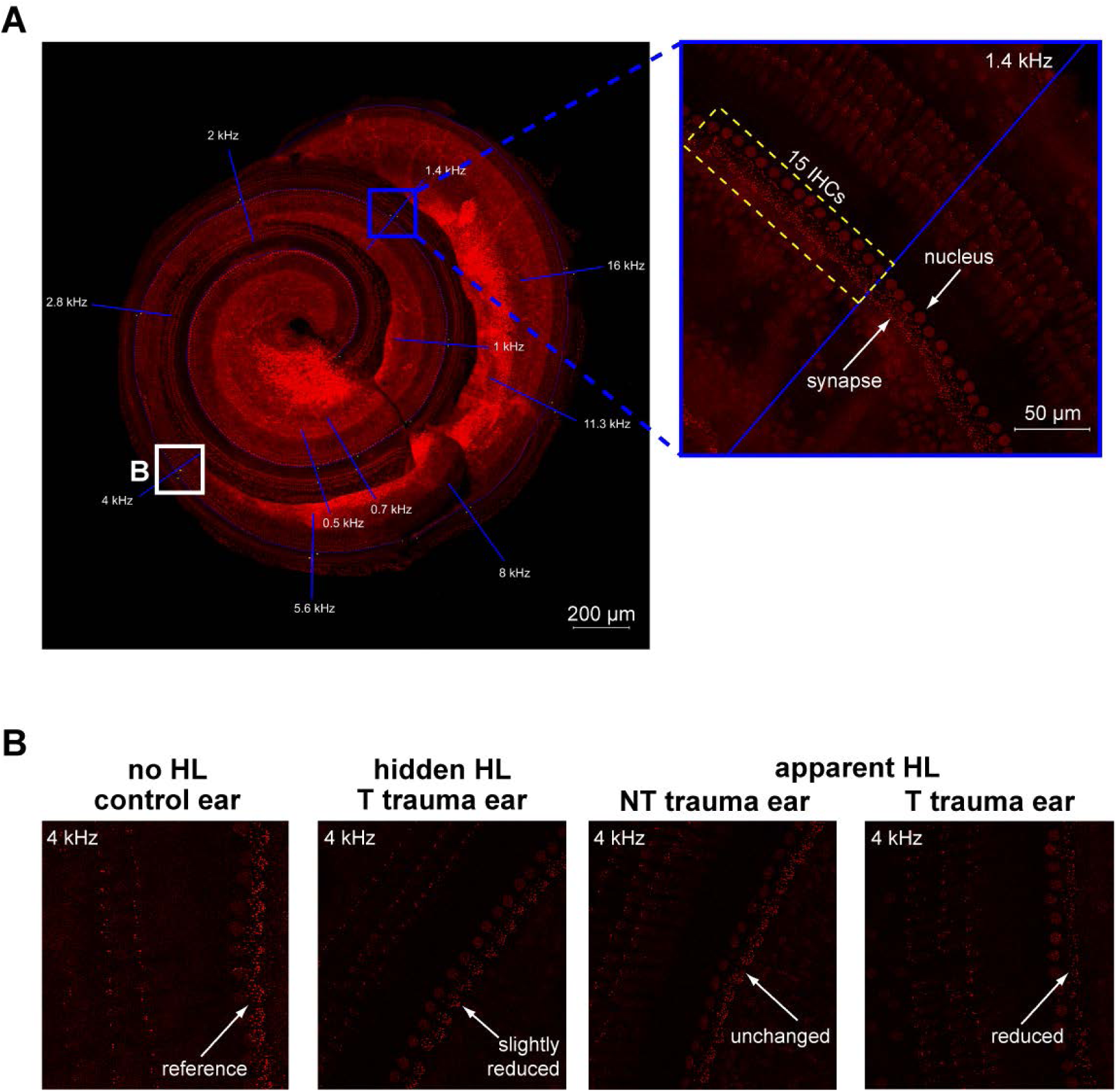
Immunohistological stainings of the Mongolian gerbil cochlea. **A**: Left: Overview of an isolated cochlea immunostained by CTBP2. Blue lines indicate the borders of the different half octave regions indicated. Right: Magnified image depicting the synapses of the 15 IHCs counted (yellow rectangle) directly above a frequency border. **B**: Exemplary images depicting the 4 kHz region of the cochlea (from left to right) of a control ear and the trauma sides of an animal with hidden hearing loss (HL) with behavioral signs of tinnitus, an animal with apparent hearing loss without behavioral signs of tinnitus (NT) and an animal with apparent hearing loss and tinnitus (T). Reduced synapse counts were only seen in animals with tinnitus, but not in control animals or animals with apparent HL but no tinnitus.

## Results

### Development of NIHL and tinnitus

15 animals were traumatized monaurally, 4 animals (8 ears) were used as sham control (cf. Methods). Development of a significant noise trauma induced HL (“apparent hearing loss”, cf. Methods) was checked by ABR measurements 3 to 5 days after the trauma on the trauma side, and development of behavioral signs of a tinnitus percept (tinnitus group) was tested behaviorally 13 days after trauma (GPIAS, cf. Methods) and is indicated by a significant negative GPIAS change in at least one of the 5 tested frequencies; no significant negative changes have been found in the sham group. Out of the 15 traumatized animals, 5 developed apparent HL but no signs of tinnitus (non-tinnitus group, NT), 5 developed signs of tinnitus (T) but no apparent hearing loss (= animals with hidden hearing loss as confirmed by histological analysis of synaptopathy, cf. below) and 5 animals developed both.

The T animals with apparent hearing loss (5/10) as well as the T animals with hidden hearing loss (5/10) showed one affected frequency in 3 cases and two affected frequencies in 2 cases each. In other words, the 10 T animals showed 14 tinnitus frequencies overall. An overview of the mean GPIAS change (± standard deviation) for the 10 T (red symbols, orange bars: number of significant tinnitus tests as a function of test frequency,), the 5 NT animals (blue) and the 4 sham control animals is given in Figure 2.

The quantification of the noise induced HL was based on comparisons of 3 to 5 days post trauma versus pre trauma ABR based audiograms (Fig. 3A). Given are the interaction plots with F-statistics of the 2-factorial ANOVAs (factors *stimulation frequency* and *investigated groups pre and post trauma*) of the different analyses in Figs 3A, C, D and F. In no case a significant interaction was found, indicating no frequency specific but – if applicable – rather general threshold changes of the affected ears after the trauma. In the analysis shown in Fig 3A, the corresponding 1-factorial part of factor *group pre and post trauma* showed a significant effect (F(3, 552)=6.66, p<0.001) and the Tukey post-hoc tests revealed that only the mean threshold of trauma side post measurements was elevated compared to the three other conditions (p-values between p=0.03 and p<0.001). The mean ABR threshold increases (= hearing loss, HL) of the trauma ears was significant higher (Students t-test, p<0.001) compared to the control ears of the same 15 animals (Fig. 3B). No threshold shifts were seen in the animals with sham trauma (Fig. 3C), which is made clear by the corresponding 1-factorial part of factor *group pre and post trauma* showing no significant differences (F(1, 90)=2.07, p=0.16).

**Figure 2.**
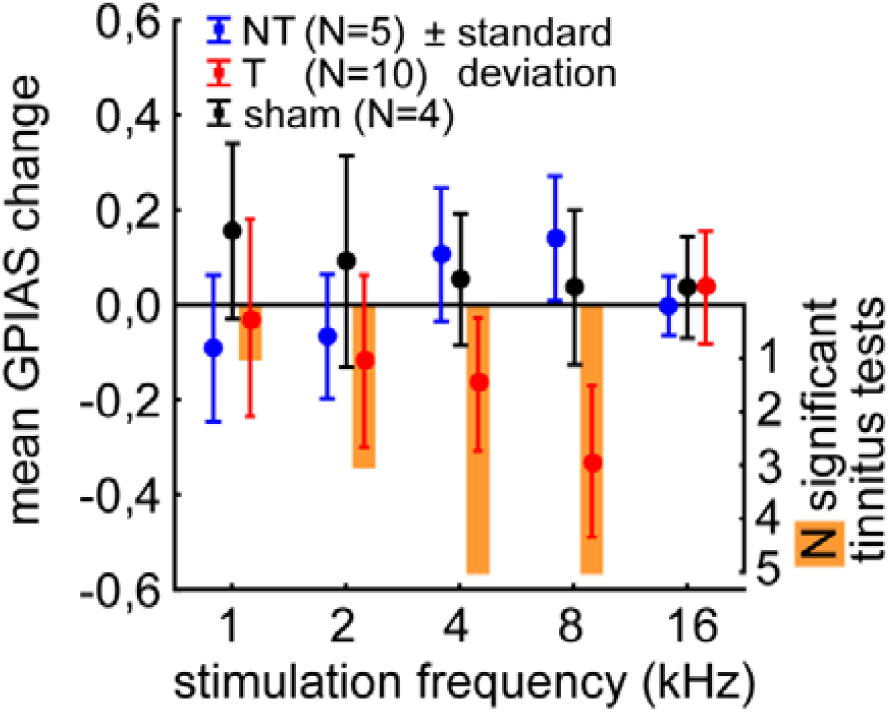
Overview of the median PPI changes in the GPIAS test for tinnitus across all 19 animals for the five frequencies tested. Blue symbols depict the mean GPIAS changes (± standard deviation) of the NT animals (no significant change), red symbol depict the changes in T animals (at least one frequency significantly affected), black symbols depict sham control results. The orange bars depict the number of significant GPAIS changes indicating a possible tinnitus percept for each frequency.

**Figure 3.**
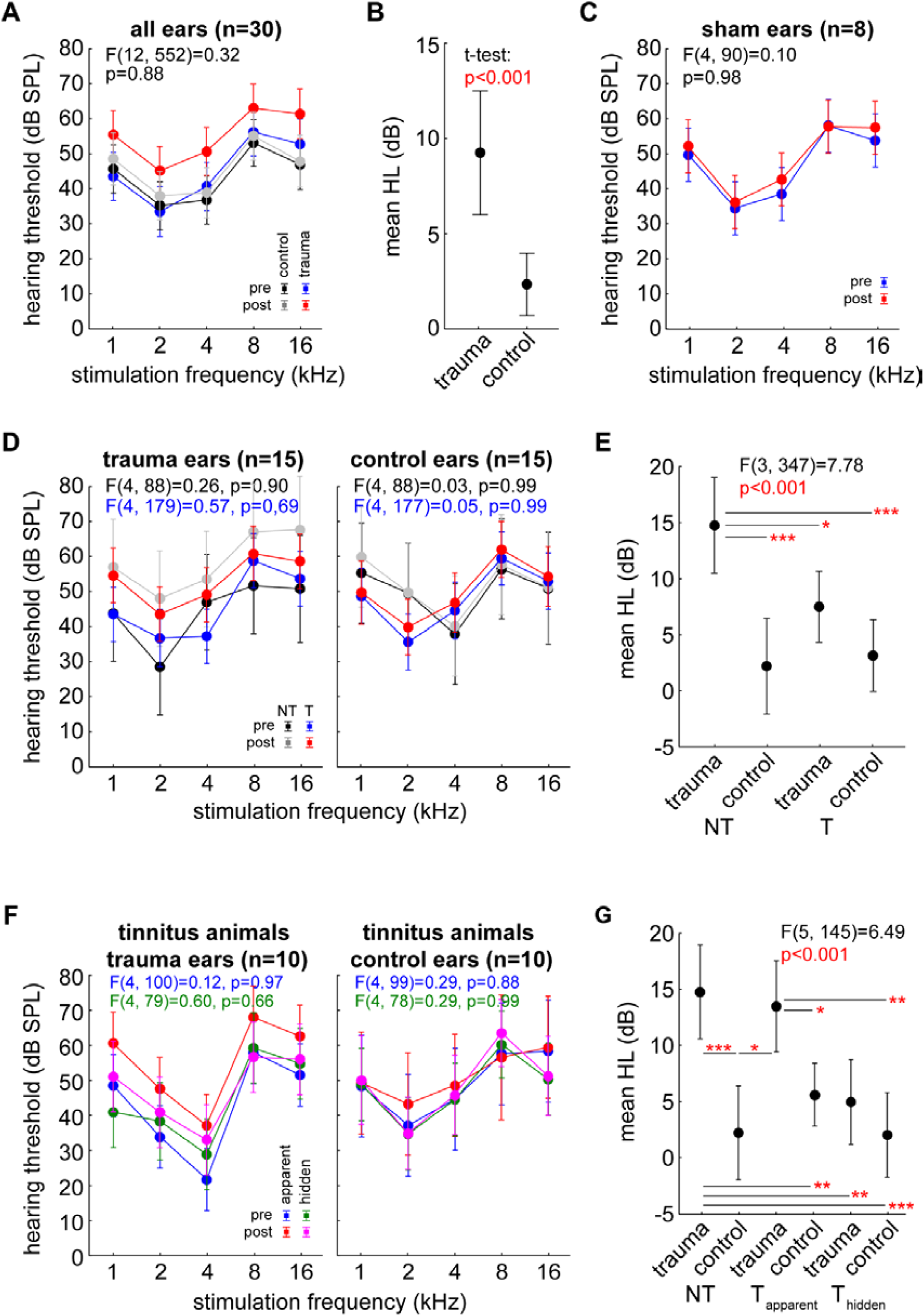
Analysis of mean hearing thresholds and mean hearing loss. **A**: Interaction plot of 2-factorial ANOVA of mean hearing thresholds (± 95% confidence interval) over the different stimulation frequencies of all animals’ (N=15) traumatized (blue and red symbols) and non-traumatized control ears (black and grey symbols) before and after trauma (number of ears: n=30). **B**: Averaged ABR threshold increase (hearing loss, HL) across all stimulation frequencies for the different ears tested by Students t-test. **C**: Interaction plot of 2-factorial ANOVA of mean pre and post treatment ABR thresholds of the 4 sham animals. **D**: Overview of mean hearing thresholds of trauma (left) and control (right) ears of the NT (black and grey symbols) and T animals (blue and red symbols). Analyses performed by four 2-factorial ANOVAs, color of F-statistics refer to corresponding interaction plot: black for NT, blue for T animals. **E**: 1-factorial ANOVA for the averaged HL data of NT and T animals. Asterisks indicate significance levels of Tukey post-hoc tests: * p<0.05, *** p<0.001. **F**: Overview of mean hearing thresholds of trauma (left) and control (right) ears of the two tinnitus animal groups with hidden (green and magenta symbols) and apparent (blue and red symbols) HL before and after trauma. Analyses performed by four 2-factorial ANOVAs, color of F-statistics refer to corresponding interaction plot: blue for T_apparent_, green for T_hidden_ animals. **G**: 1-factorial ANOVA for the averaged HL data of NT and both T groups. Asterisks indicate significance levels of Tukey post-hoc tests: * p<0.05, ** p<0.01, *** p<0.001.

As described above, we classified the animals based on, first, the individual behavior in the GPIAS test into NT and T animals (Fig. 3 D and E) and, second, based on ABR data into animals with and without a significant HL (i.e. apparent and hidden hearing loss, respectively; Fig. 3 F and G). Note that all animals in our sample without a significant noise induced HL showed synaptopathy (cf. below) and therefore are classified as “hidden hearing loss” and not “no hearing loss”. In the four 2-factorial ANOVAs (NT trauma side, T trauma side, NT control side, T control side) performed and given in Fig 3D, the mean thresholds on the trauma sides of NT (F(1, 88)=10.66, p=0.002) and T animals (F(1, 179)=8.75, p=0.004) were elevated after trauma. This was not the case on the control sides (NT: F(1, 88)=0.14, p=0.70; T: F(1, 177)=0.79, p=0.38). NT animals showed in a 1-factorial ANOVA an apparent HL on the trauma side (Fig. 3E) that was significantly higher than the threshold change on the control side (Tukey post-hoc test, p<0.001) and also different from the control side (Tukey post-hoc test, p<0.001) and trauma side (Tukey post-hoc test, p<0.05) of all T animals. The HL of the trauma sides of all T animals did not show a significant difference to their control sides (Tukey post-hoc test, p=0.22). Separating the T animals into those with apparent and hidden hearing loss (Fig. 3F) and analyzing the data by four 2-factorial ANOVAs (cf. above), only in the apparent HL T animals a significant mean threshold shift on the trauma side could be identified (F(1, 100)=17.87, p<0.001). In a final step, the HL of all 15 animals was investigated in one single 1-factorial ANOVA with the groups NT, T_apparent_ HL and T_hidden_ HL for trauma and control ears (Fig. 3G). NT (Tukey post-hoc test, p<0.001) and T_apparent_ (Tukey post-hoc test, p<0.05) animals did show HL on the trauma sides compared to their control sides. Neither trauma nor control sides were significantly different between these groups. T_hidden_ animals did not show a significant HL at all on their trauma side when compared to their control side or any other control side. Taken together these findings indicate: The amount of ABR threshold shift may not be the exclusive cause for the development of a tinnitus percept, as it may arise also without (ABR-) detectable hearing loss.

### Effects of noise trauma on ABR and their relation to tinnitus development

The peak-to-peak amplitudes of ABR waves I&II, III, IV and V obtained at 65 dB SPL were analyzed in detail by 2-factorial ANOVAs. We compared the pre and post trauma data of both ears for all animals with apparent HL (T, NT) in one analysis. Hidden HL T animals were investigated separately. Generally, the ABR amplitude data of the animals’ affected and control ears were not significantly different (exception NT animals wave I&II, cf. below) but especially for the earlier waves only relatively few valid wave amplitudes could be detected on the affected ear which may reduce the strength of the analysis. Nevertheless, for the apparent HL groups, the Tukey post-hoc tests revealed decreases in NT animals’ wave I&II on the trauma side only, as well as wave V peak-to-peak amplitudes in both ears after trauma (Fig. 4, upper left panel, lower right panel). In T animals with hidden HL no significant differences could be found.

**Figure 4.**
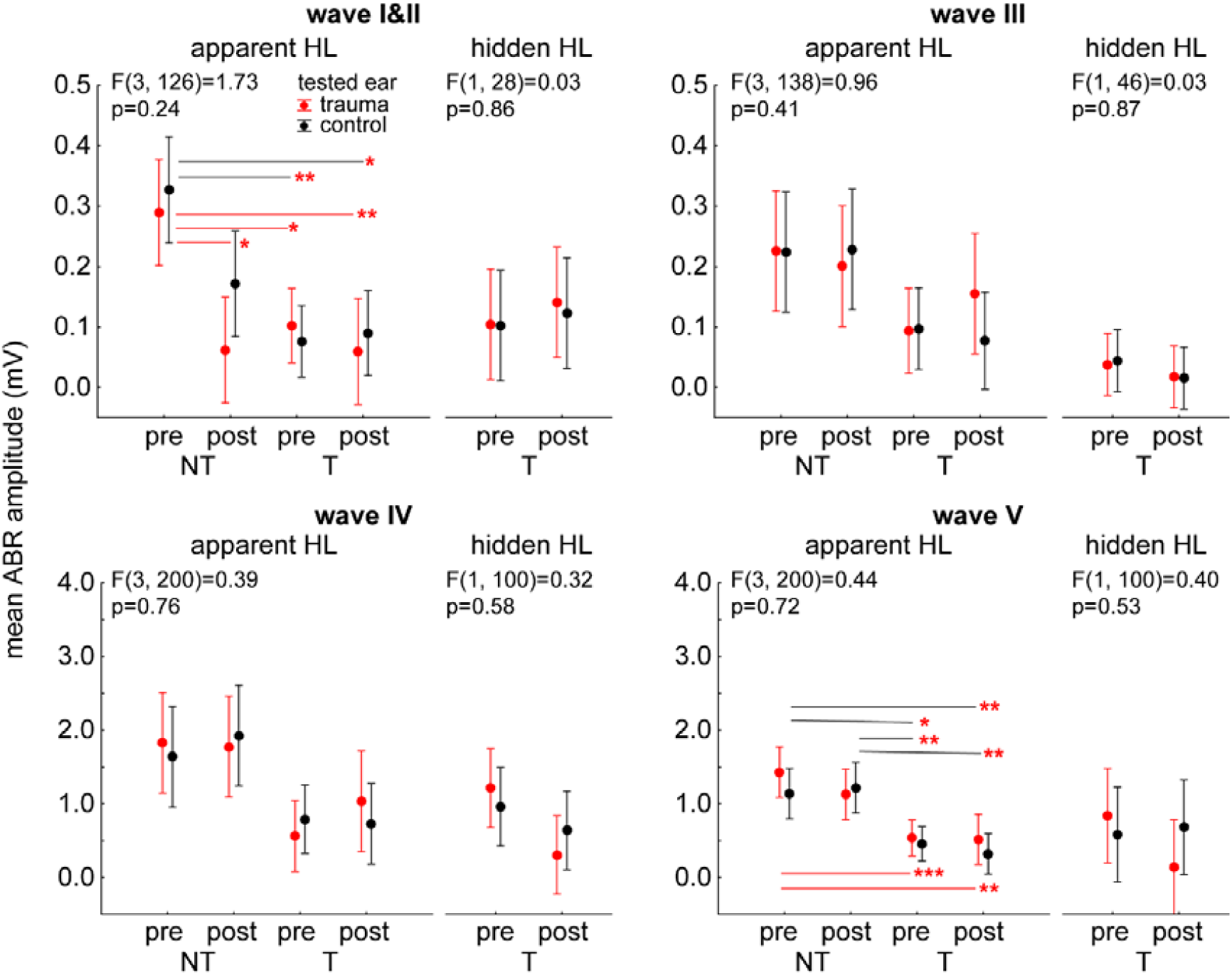
Overview of ABR wave analyses performed with 2-factorial ANOVAs. Interaction plots of mean peak-to-peak amplitudes of both ears of single ABR waves at a stimulation intensity of 65 dB SPL (panels) in animals with apparent HL averaged across all stimulation frequencies at each panels left side; on panels’ right sides the corresponding plots of animals with hidden HL are displayed. Asterisks indicate significance values of the Tukey post-hoc tests; * p<0.05, ** p<0.01, *** p<0.001.

Note that amplitudes of NT animals – in line with our earlier results [15, 16] – were significantly larger than T animals’ amplitudes in all waves (I&II: F(3, 126)=13.63, p<0.001; III: F(3, 138)=5.00, p=0.003; IV: F(3, 200)=7.78, p<0.001; V: F(3, 200)=16.08, p<0.001) while the two T groups showed similar amplitudes (four separate 1-factorial ANOVAs, always p>0.05).

In a further analysis (Fig. 5), we investigated the 90 dB SPL ABR responses (valid detection of waves in all cases) and pooled the data from both ears of each T animal (apparent HL with tinnitus and hidden HL with tinnitus) as nearly no difference in ABR was found between ears and tinnitus animal groups (cf. above). We compared *pre and post trauma* ABR RMS data aligned to the lowest tinnitus frequency determined with the behavioral GPIAS test. The reasoning behind this approach was that due to the differences in perceived tinnitus frequencies (cf. Fig. 2) any possible specific tinnitus related amplitude effect may be blurred. By such re-aligning to the individual tinnitus frequency, we found a significant general effect of a trauma induced decrease of overall ABR RMS values (Fig. 5, left panel). Interestingly, an ‘edge’ effect was observed for responses below the tinnitus frequency compared to the ABR RMS values at and above the tinnitus frequency (Fig. 5, center panel). In the significant interaction of both factors (*distance from tinnitus frequency* and *time point*) we found significant drops in response amplitudes after trauma at the tinnitus frequency and one octave below this frequency (Tukey post-hoc tests, Fig. 5, right panel) which indicated a specific trauma effect on the ABR at these frequencies in T animals.

**Figure 5.**
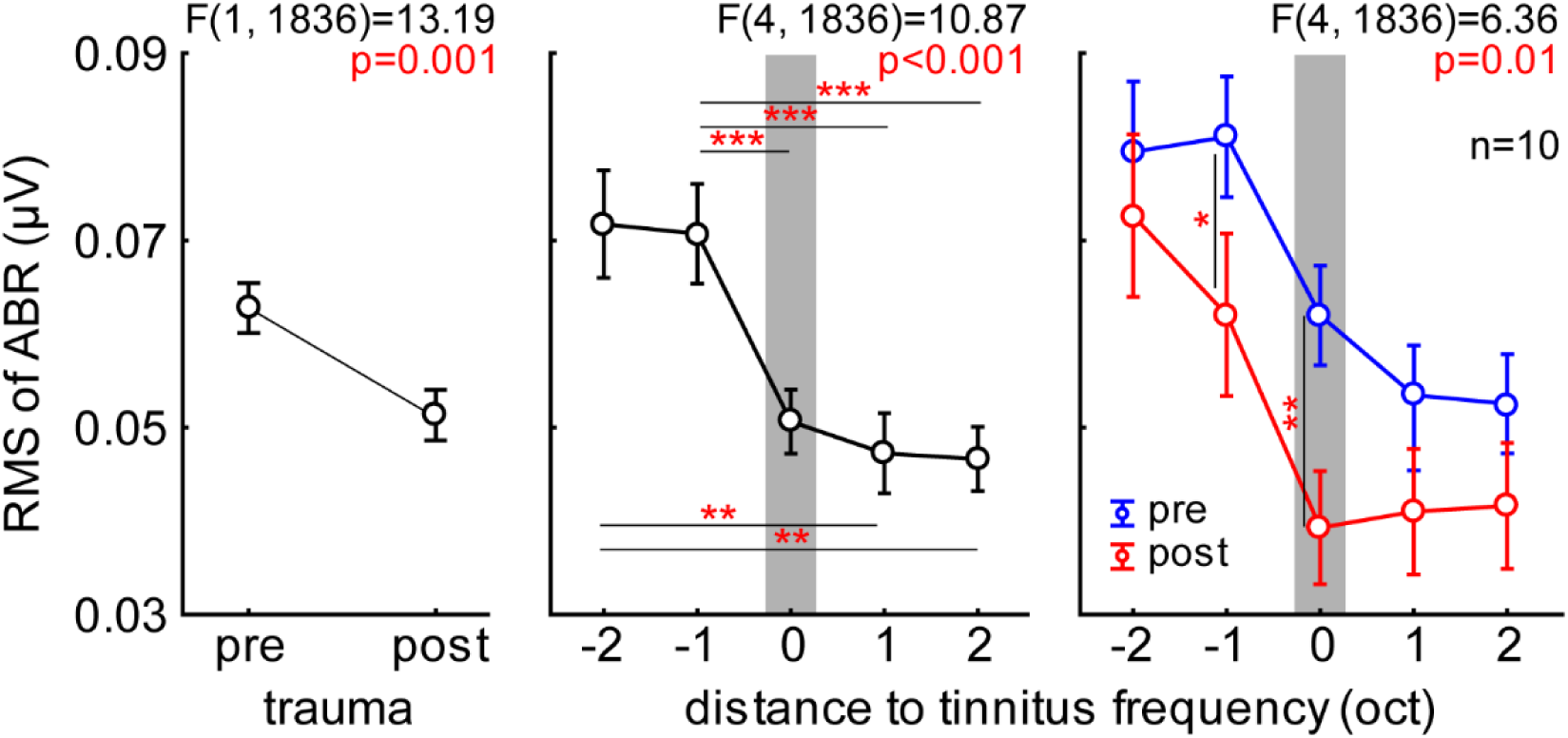
Overview of 90 dB SPL ABR RMS 2-factorial ANOVA analysis of both T groups. RMS values of the ABR signals averaged across all frequencies, intensities and both ears as a function of distance from the individual tinnitus frequency and time point of measurement. The vertical grey bars indicate the tinnitus frequency the data are aligned to. Asterisks indicate significance values of the Tukey post-hoc tests; * p<0.05, ** p<0.01, *** p<0.001.

### Synapse counts of IHC afferents

We compared the IHC afferent synapse counts of the control ears and the traumatized ears separately (Fig. 6A, B) as well as the sham control ears (Fig. 6C). We found that a significant loss of synapses could be only found at frequency locations above the trauma frequency (Fig. 6B, Mann-Whitney U-test, p=0.005), and sham ears did not show such difference (synapse count difference of left vs. right sham ear, tested: frequencies up to 2 kHz vs. frequencies above 2 kHz; Mann-Whitney U-test, p>0.05). Most importantly, all T animals – independent of the type of HL - showed a significant loss of synapses in the traumatized ear only. This is obvious from, first, separated synapse counts of NT and T animals (Fig. 6D) and the difference between individual control and trauma ears of these two groups for cochlear frequency locations up to the trauma frequency and above that frequency (Fig. 6E). In contrast, we found no significant difference between both animal groups for the low frequency regions (median NT (lower, upper quartile): NT: 0.92 (0.33, 1.67), T: 0.10 (−1.42, 2.75), Mann-Whitney U-test, p>0.05) but a significant decrease of synapses in the T animals compared to the NT animals (NT: 0.93 (−1.00, 2.00), T: -1.99 (−5.10, 0.24), Mann-Whitney U-test, p<0.001) in the higher frequency range above the trauma. The effect of the type of the HL can be seen in a second analysis in which we specifically looked at the synapse change in the T animals but separated the data of the animals with hidden and apparent HL (Fig. 6F). When comparing the synapse loss data dependent on behavior and type of HL (Fig. 6G) we found for lower frequency locations up to the trauma frequency no significant differences between the three animal groups (NT: 0.92 (0.33, 1.67), T_hidden_: 0.79 (−1.42, 3.11), T_apparent_: -0.35 (−2.12, 1.10), Kruskal-Wallis ANOVA: H(2, 56)=4.51, p=0.10). Above the trauma we found significant differences between the three groups (NT: 0.93 (−1.00, 2.00), T_hidden_: -3.07 (−4.02, -1.86), T_apparent_: -5.17 (−8.97, - 1.98), Kruskal-Wallis ANOVA: H(2, 99) =30.61, p<0.001). Both T animal groups did show significantly lower synapse numbers compared to NT animals (multiple comparison of mean ranks post-hoc tests: NT vs. T_hidden_: p<0.001, NT vs. T_apparent_: p<0.001). Note that the NT animals all showed apparent HL in their audiograms. Additionally, the animals with tinnitus and apparent HL showed even stronger synapse loss than the T animals with hidden HL (T_hidden_ vs. T_apparent_: p=0.04).

**Figure 6.**
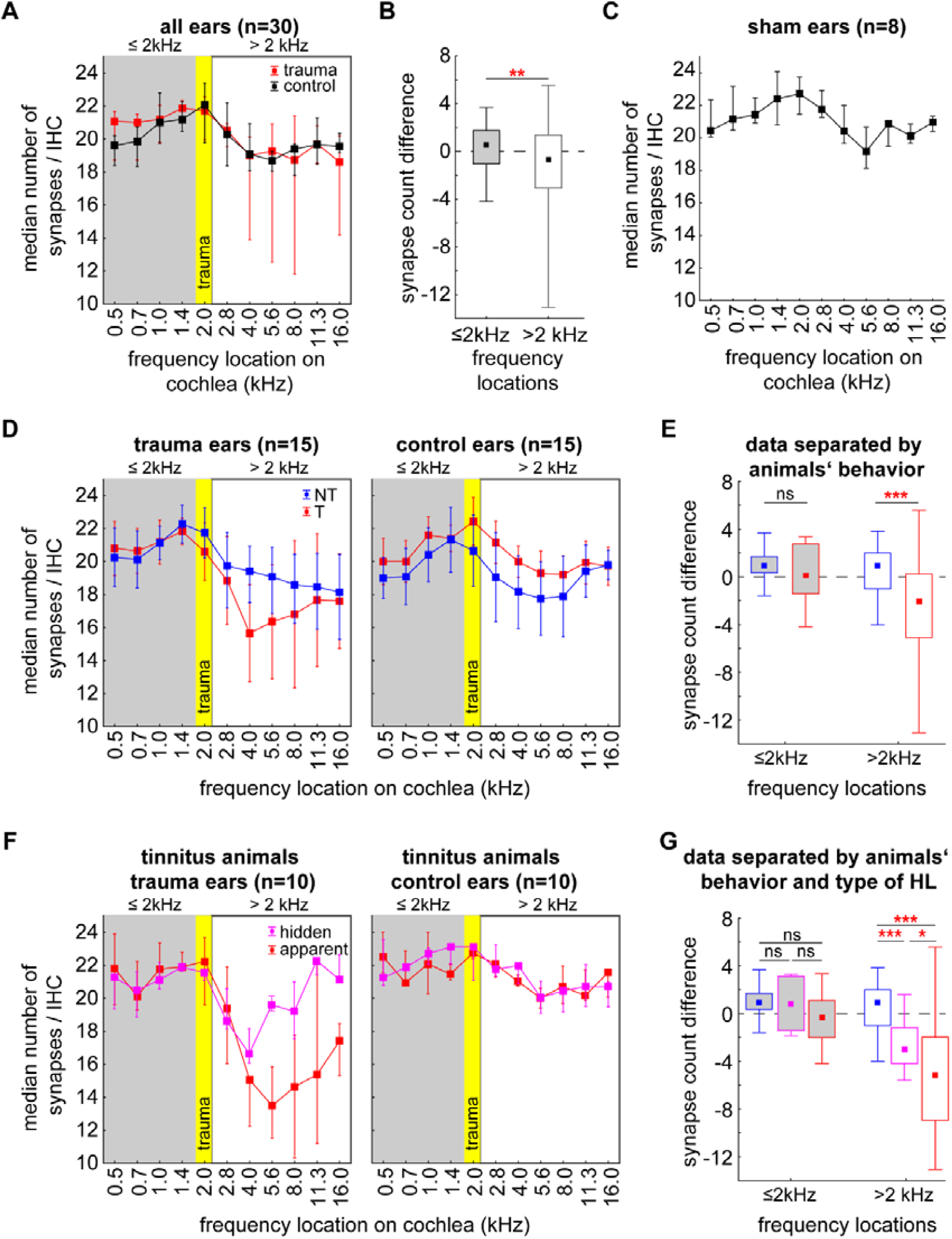
IHC synapse counts in animals with and without behavioral signs of tinnitus assessed by GPIAS testing using Mann-Whitney U-tests or Kruskal-Wallis ANOVAs. **A**: Median synapse/IHC counts as a function of cochlear location of trauma and control cochleae of all animals. Yellow box: trauma frequency; grey area: frequency locations below trauma; white area: frequency locations above trauma. **B**: Median synapse count differences between individual control and trauma ears separated for frequency regions at and below the trauma frequency and above that frequency. Asterisks show significance level of Mann-Whitney U-test: ** p<0.01. **C**: Median synapse/IHC counts as a function of cochlear location of sham ears. **D**: Median synapse/IHC counts as a function of cochlear location of trauma (left panel) and control (right panel) cochleae separated by GPIAS tinnitus assessment (NT: non-tinnitus, blue symbols; T: tinnitus, red symbols). **E**: Median synapse count differences between individual control and trauma ears of NT and T animals separated for frequency regions at and below the trauma frequency and above that frequency. Asterisks show significance level of Mann-Whitney U-test: *** p<0.001. **F**: Median synapse/IHC counts as a function of cochlear location of trauma and control cochleae for T animals only, separated by HL group (hidden HL: magenta symbols; apparent HL: red symbols). **G**: Median synapse count differences between individual control and trauma ears of NT and T_hidden_ and T_apparent_ separated for frequency regions at and below the trauma frequency (Kruskal-Wallis ANOVA: H(2, 56)=4.51, p=0.10) and above that frequency (Kruskal-Wallis ANOVA: H(2, 99) =30.61, p<0.001). Asterisks show significance level of multiple comparison of mean ranks post-hoc tests: ns not significant, * p<0.05, ** p<0.01, *** p<0.001.

## Discussion

In this study we investigated the synapse loss at IHCs after a mild acoustic trauma – previously shown in mice (Liberman *et al*., 2015) – in the Mongolian gerbil and relate it to possible behavioral signs of tinnitus (Turner, 2007; Ahlf *et al*., 2012). Further, we compared animals with hidden and apparent HL and investigated the relationship between IHC synaptopathy, type of HL and tinnitus development. For noise trauma induction, we used a well-established method in our lab, consisting of a 2 kHz pure tone, thereby targeting only a specific region of the cochlea. A specific goal was to evaluate, if the extend of synapse loss within the cochlea is correlated to tinnitus-related, maladaptive processes along the auditory pathway as described earlier (Ruttiger *et al*., 2013; Hickox & Liberman, 2014). The main finding reported here is a frequency specific reduction in IHC synapse counts, most prominently one octave above the trauma frequency, only in animals with behavioral signs of tinnitus and independent of the kind of HL, hidden or apparent.

### Methodological considerations of the behavioral paradigm

For identification of behavioral signs of tinnitus in our animal model (the Mongolian gerbil), we use a gap-startle paradigm adopted from Turner and colleagues (2006), but with improved statistics (Schilling *et al*., 2017). The measurement of the PPI of the ASR 13 days post trauma relative to pre trauma conditions was analyzed (Tziridis *et al*., 2012). This approach has the advantage of being independent of training (e.g., Jastreboff *et al*., 1988a; Jastreboff *et al*., 1988b; Ruttiger *et al*., 2013) but is considered not to be failsafe (for review see Eggermont, 2013). To overcome this problem we always relate the behaviorally determined classification of tinnitus and non-tinnitus animals with additional data, like electrophysiological recordings (Ahlf *et al*., 2012; Tziridis *et al*., 2014; Tziridis *et al*., 2015) or in this report electrophysiological ABR recordings and histological data. Nevertheless, one has to be critical of the method, as even with the additional data, we can only speak of correlations of behavior and other data in this animal model. A synaptopathy in different rodent models with or without tinnitus development (for review see Liberman & Kujawa, 2017; Heeringa *et al*., 2018; Pienkowski, 2018) using this behavioral paradigm have been described. One may speculate about possible reasons for the differences in pathology, e.g., different individual stress levels, unintentional additional noise exposure or unnoticed peripheral damage (Longenecker & Galazyuk, 2016) or a damage to the outer hair cells (OHC, cf. below). Nevertheless, using the described approach in overall 19 animals (15 traumatized animals plus 4 sham controls), we found a tinnitus percept in hidden (n=5) or apparent hearing loss (n=5) animals (10/15, 66.7%) and no behavioral signs of tinnitus in 5 animals (33.3%). This is well in the range of our previous results with a tinnitus developing in 65% to 80% of animals with NIHL (Ahlf *et al*., 2012; Tziridis *et al*., 2014; Krauss *et al*., 2016a) even though the major differences to our usual approach was the use of monaural rather than binaural traumata for obtaining intra-individual controls.

### Noise trauma induced tinnitus – predisposition and development

The differences in ABR amplitudes between T and NT animals, with NT animals showing generally larger amplitudes than T animals even before induction of noise trauma, are consistent with our earlier results (Tziridis *et al*., 2015). We were able to reproduce the generally larger peak-to-peak ABR amplitudes in all identified wave complexes in NT compared to T animals. In our earlier studies we compared ABR before and immediately after binaural noise trauma and found no differences between pre and post waves with the exception of a general increase of wave V amplitudes (inferior colliculus) in T animals which was usually stable up to 7 days after the trauma. Here we compared the pre trauma ABR waves with those obtained 3 to 5 days after monaural trauma, that is, probably within the time frame of the chronic manifestation of a tinnitus percept (cf. Ahlf et al., 2012) which we here measured 13 days after the trauma. As the response levels of T animals are significantly smaller than those of NT animals and especially earlier waves up to wave III of the trauma ear were often hard to detect at 65 dB SPL, a potential difference between control and trauma side might be hidden. Nevertheless, at least for NT animals no significant changes of wave I&II were found on the control side, while the amplitudes on the trauma side were significantly reduced (Fig. 4, upper left panel). This may indicate either a decrease of neuronal activity (e.g., Sergeyenko *et al*., 2013) or a loss of synchronization (cf. Berlin *et al*., 2003) in the early auditory signals of these animals and it resembles our earlier data on a predisposition for tinnitus development (Ahlf *et al*., 2012; Tziridis *et al*., 2015). But as we found no other differences between corresponding trauma and control ears this could also indicate that the plugging did not completely protect the control ear and by that activity of both sides was similar.

Independent of the individual waves at 65 dB SPL, the more general RMS values at 90 dB SPL (including all five waves) of the animals with behavioral correlates of a tinnitus percept (Fig. 5) clearly showed a significant decrease of neuronal activity and/or synchronization after the trauma, especially at the tinnitus frequency and one octave below. This indicates, first, that the trauma itself worked also in the T animals. And second, that around the tinnitus frequency neuronal (mal-) adaptation seems to be strongest. This is in line with several models of tinnitus development (Krauss *et al*., 2019) and may further strengthen our argument, that the behavioral measurement of tinnitus with the GPIAS paradigm is relying on a real percept.

### Synaptopathy and tinnitus

We found significant synaptopathic effects at the IHC of the cochlea in all animals with behavioral signs of tinnitus, irrespective of the type of HL, hidden or apparent. Interestingly, the strongest reduction of synapses (roughly 25%) was not located at the trauma frequency but about one octave above (cf. Fig. 6), consistent with effects reported in mice (Hickox & Liberman, 2014; Liberman *et al*., 2015) and rats (Ruttiger *et al*., 2013). In addition, this frequency range has frequently been reported to reflect the perceived tinnitus frequency in Mongolian gerbils (e.g., Nowotny *et al*., 2011). No such loss of IHC synapses was found in NT animals even though they all showed apparent HL at the level of the hearing thresholds not different from the one in tinnitus animals with apparent HL (cf. Fig. 3). It is notable that the mild hearing loss in our study is not accompanied by a loss of IHCs, which is in line with an earlier study demonstrating that apoptosis is only found after strong traumata with threshold increases of at least 50 dB (e.g., Chen & Fechter, 2003). The exact cause of the finding of apparent hearing loss without synaptic loss in these animals remains elusive, but a number of possibilities are conceivable, like stereocilia damage (Engstrom, 1983; Liberman & Kiang, 1984) to high threshold hair cells (e.g., Wang *et al*., 1997) or the animals’ noise protection efferent system (e.g., Zheng *et al*., 1997a; Zheng *et al*., 1997b; Kirk & Smith, 2003; Maison *et al*., 2013) being more efficient. The possibility that such synapse reductions may have been overlooked is very unlikely as we scanned the cochlea with relatively high spectral resolution, i.e. in half octave steps (cf. Methods). Therefore, the apparent HL, i.e. increased hearing thresholds, in our NT group do not seem to be based on IHC synapse damage but next to the possibilities described above maybe due to OHC damage. Support to the idea of OHC damage leading to hidden HL also comes from our hidden HL T animals, as their overall loss of synapses is significantly lower than in T animals with apparent hearing loss. We conclude from these observations that tinnitus development (or the lack of it) seems to depend on afferent IHC innervation. This conclusion is in line with our model of tinnitus development, where a reduction of cochlear input into the auditory system is the trigger for compensatory mechanisms within the auditory system that constantly optimize information transmission between the cochlea and the auditory system by means of stochastic resonance (Krauss *et al*., 2016b; Krauss *et al*., 2018; Krauss *et al*., 2019). Additionally, mild traumata seem to lead to a higher percentage of animals with behavioral signs of tinnitus than more severe traumata (Devarajan *et al*., 2012; Kiefer *et al*., 2015), which may correlate with the variable proportion of damage to IHC and OHC, respectively.

Taken together, we conclude that at least in our animal model, a hearing threshold increase induced by mild acoustic trauma is associated with synaptopathy at the IHC and/or OHC damage (Dallos & Harris, 1978; Harrison, 1981; Schaette & Kempter, 2006), while the tinnitus percept itself seems to be associated primarily with IHC synaptopathy, independent of a threshold increase.

## Acknowledgements

The present work was performed in fulfillment of the requirements for obtaining the degree „Dr. med. dent.” at the Friedrich-Alexander University Erlangen-Nürnberg (FAU). We are especially grateful to Dr. Tobias Moser, Georg-August-University Göttingen, for methodical support for the whole mount cochlea preparation.

## Author Contributions

JF and IBD performed the animal experiments ES and OW performed the whole mount histology JF performed the microscopy KT performed the statistical analysis PK and AS provided valuable input for data interpretation and manuscript discussion KT and HS wrote the manuscript

## Data Accessibility Statement

Data and python code will be made accessible for every interested scientist upon personal request.

ABR: auditory brainstem response
ASR: acoustic startle response
GPIAS: gap prepulse inhibition of acoustic startle
HL: hearing loss
IHC: inner hair cells
NT: animals without behavioral signs of tinnitus = non-tinnitus animals
OHC: outer hair cells
PPI: prepulse inhibition
T: animals with behavioral signs of tinnitus = tinnitus animals

